# DuplexFM: Transferable small-RNA target representations link miRNA interactions to siRNA efficacy prediction

**DOI:** 10.64898/2026.08.06.743413

**Authors:** Baiming Chen, Jiaqi Yin, Fei Jia, Mingjun Yang

## Abstract

MicroRNAs (miRNAs) and small interfering RNAs (siRNAs) share Argonaute-mediated guide-target recognition, yet quantitative siRNA efficacy measurements are substantially scarcer and more costly to generate than miRNA-target interaction data. We therefore asked whether miRNA interaction data could provide transferable supervision for siRNA efficacy prediction. Here we present DuplexFM, a biologically grounded framework that uses sample-specific gates to integrate five evidence sources: pairing and sequence-context priors, duplex energetics, experimentally supervised mRNA accessibility, target-to-guide cross-attention, and contextual token-pair compatibility. The accessibility expert, trained on nucleotide-resolution icSHAPE measurements, achieved a held-out nucleotide-level Pearson correlation of 0.627 and evaluated accessibility at seed match and energy-supported candidate sites. On miRBench v7, three independently trained DuplexFM models achieved a macro APS of 0.873±0.002, soft-voting increased this to 0.876 and yielded the highest APS on all four test sets. We then froze the miRNA-trained representation and trained only a lightweight residual head with 24 siRNA-specific descriptors. Transfer improved Pearson and Spearman correlations, AUPRC, and F1 over the descriptor-only baseline in all six evaluation settings. The ensemble achieved the highest Pearson and Spearman correlations in four settings, whereas OligoFormer remained stronger on Huesken and Takayuki. These findings show that experimentally grounded accessibility and miRNA-derived interaction representations provide complementary, transferable information, supporting a parameter-efficient route towards unified modeling of Argonaute-guided RNA regulation. Code and data are available at https://github.com/cbaiming/DuplexFM.

## 1 Introduction

MicroRNAs (miRNAs) are small non-coding RNAs that post-transcriptionally regulate gene expression by binding to target transcripts, typically within 3’ untranslated regions (3’ UTRs), and promoting translational repression or transcript degradation [1, 2]. Small interfering RNAs (siRNAs) are another class of small non-coding RNAs that mediate sequence-specific gene silencing by guiding the degradation of complementary target mRNAs [3]. miRNA-and siRNA-mediated regulation are involved in Cell development, differentiation, tumorigenesis, immune responses, and many other biological processes [4, 5, 6, 7, 8, 9]. Although miRNAs and siRNAs differ in biogenesis and canonical targeting patterns, they act through closely related RNA-induced silencing complex (RISC)-mediated mechanisms [10]. For instance, siRNA off-target effects arise from miRNA-like repression, leading to unintended regulation of transcripts, often via target sites located in 3’ untranslated regions (UTRs) [11]. This mechanistic overlap raises the possibility that representations learned from miRNA targeting may provide transferable information for siRNA efficacy prediction.

This transfer learning strategy is particularly attractive because the two tasks differ markedly in the amount of available labeled data. High-throughput Argonaute (AGO) ligation assays and chimeric eCLIP experiments, together with systematic data curation, have expanded sequence-resolved miRNA–target resources to encompass millions of training pairs. For instance, the miRBench collection Manakov2022 [12, 13] contains over two million miRNA binding-site entries. These miRNA–target sequences serve as a rich source of information for learning or computing pairing preferences, duplex energetics, local target accessibility, and regional sequence context. In contrast, obtaining high-quality labels for siRNA efficacy requires controlled perturbation experiments, and existing data remain scattered across relatively small, heterogeneous studies. For example, the OligoFormer study aggregated 9 published datasets yet assembled only 3,714 siRNAs targeting 75 mRNAs, explicitly identifying the limited size and representational diversity of training data as a constraint on model generalization [14]. This data asymmetry motivates learning a general representation of small-RNA target interactions from the abundant miRNA resource and transferring it to the data-scarce siRNA efficacy task.

Over the past two decades, a large number of computational models have been developed for miRNA-mRNA interaction prediction, evolving from rule-based and thermodynamics-driven methods to machine learning and deep learning frameworks. Early representative tools, such as miRanda, align miRNA sequences to candidate target regions using weighted sequence complementarity, duplex free energy, and cross-species conservation, and were widely used for genome-scale scanning of candidate sites [15]. TargetScan places stronger emphasis on canonical seed pairing, evolutionary conservation, and site context. Latest versions further introduced context/context++ scoring, which combines site type with multiple contextual features, including local AU composition, 3’-supplementary/compensatory pairing, etc. [2, 16]. More generally, traditional sequence-based predictors usually rely on a combination of several biologically motivated features, including seed-match type, thermodynamic stability, evolutionary conservation, and local structural accessibility [17], but their performance is limited by incomplete modeling of non-canonical interactions and by insufficient incorporation of cellular and transcriptomic context, although many strongly functional sites in mammalian systems remain canonical [18]. To address these limitations, machine learning models were introduced to learn predictive rules from experimentally supported interactions. Representative examples include mirSVR [19], which uses support vector regression to rank miRanda-derived candidate sites by expected repression strength, as well as classifiers such as the SVM-based MultiMiTar [20] and the random-forest-based TarPmiR [21], both of which rely on engineered sequence and contextual features. More recently, deep learning methods have become increasingly prominent because they can learn task-relevant representations directly from sequence inputs, reducing reliance on manually engineered features. For example, miRAW uses deep learning on raw sequence inputs and is designed to recognize both canonical and non-canonical target patterns without relying on predefined descriptors [22]. miTAR combines convolutional and recurrent neural networks to jointly model spatial and sequential characteristics of miRNA– target pairs [23]. TargetNet employs deep residual neural networks to predict functional target sites with relaxed candidate-site selection criteria [24]. More recent models, such as TEC-miTarget, extend deep sequence learning to both sequence- and transcript-level prediction by aggregating candidate-site information within transcripts [25]. Beyond sequence-centered modeling, recent work has also explored knowledge-guided prioritization frameworks. For example, miRTarDS uses Sentence-BERT-derived semantic similarity to model disease association between miRNAs and target genes, showing that disease knowledge complements sequence evidence in functional MTI prioritization [26].

The computational prediction of siRNA-mRNA efficiency has also evolved from empirical sequence rules and thermodynamically based heuristic methods to statistical learning and deep learning frameworks. Early studies, including Reynolds rules [27] and Ui-Tei guidelines [28], defined the core determinants of siRNA efficacy. Tools such as siDirect [29] and OligoWalk [30] apply these principles to actual siRNA design. Subsequent models, including the neural network-based framework by Huesken et al [31]. And the explainable predictor by Vert et al. [32], marked a shift towards data-driven silent efficiency prediction. At the same time, Jackson et al. [11] recognized the widespread seed-mediated off-target silencing, promoting the development of tools such as siSPOTR [33] by taking specificity as a major consideration. Recently, methods such as OligoFormer [14] have further advanced the field by applying modern deep representation learning to siRNA design.

Despite these advancements, miRNA and siRNA prediction have often been treated as separate modeling problems. miRNA target prediction mainly focuses on partial complementary recognition, canonical and non-canonical site architectures, and transcript-level contextual determinants, while siRNA prediction is mainly driven by on-target silencing efficiency, thermodynamic asymmetry, and off-target properties [7, 28, 11]. However, these two problems are mechanistically related and partially overlapping. In both cases, small RNAs act as Argonaute-loaded guides that recognize target transcripts through sequence-directed interactions, and seed pairing plays a central role in target search and repression. Notably, many siRNA off-target effects arise through miRNA-like seed-mediated recognition [11], suggesting that the two problem domains may share transferable sequence features and targeting principles. For instance, although the TargetScan tool is a miRNA tool, it also analyses siRNA data [16]. We therefore hypothesize that a unified deep learning framework could learn shared principles of small-RNA-guided target recognition from miRNA–mRNA interaction data and transfer this knowledge to siRNA prediction.

Foundation models have opened new opportunities for learning representations directly from biological sequences and have demonstrated strong potential as general-purpose encoders for DNA [34], RNA [35, 36, 37], and proteins [38]. RNA foundation models can capture contextual sequence dependencies that are difficult to encode manually, making them attractive for downstream RNA prediction tasks. However, small RNA targeting is fundamentally a paired-sequence problem rather than a single-sequence representation problem: the relevant object is the interaction between a miRNA or siRNA guide and a local target mRNA fragment. Directly applying pretrained RNA encoders as independent feature extractors may obscure biologically interpretable signals such as seed-site structure and local AU context. Effective modeling of small RNA targeting therefore requires interaction-aware architectures that can align small RNA and target representations, distinguish informative token-level contacts from background sequence context, and preserve biologically meaningful priors.

In this work, we ask whether interaction knowledge learned from large-scale miRNA-mRNA experiments can be transferred to improve siRNA efficacy prediction, thereby complementing costly and experimentally demanding siRNA-specific measurements. To address this question, we developed DuplexFM, a multi-expert framework that combines RNA-FM [35] and mRNABERT [39] representations with five complementary signals: Bio23 pairing and sequence priors, IntaRNA21 energetic and duplex-geometry features, experimentally supervised mRNA accessibility, target-to-guide cross-attention, and context- and position-aware score-matrix attention (Fig. 1). All five expert representations are independently modulated by a sample-specific sigmoid router, with Bio23 and IntaRNA21 providing mechanistically grounded core evidence. For siRNA transfer, the miRNA-trained expert representations are frozen and coupled to a lightweight TD24-anchored residual head.

**Figure 1:**
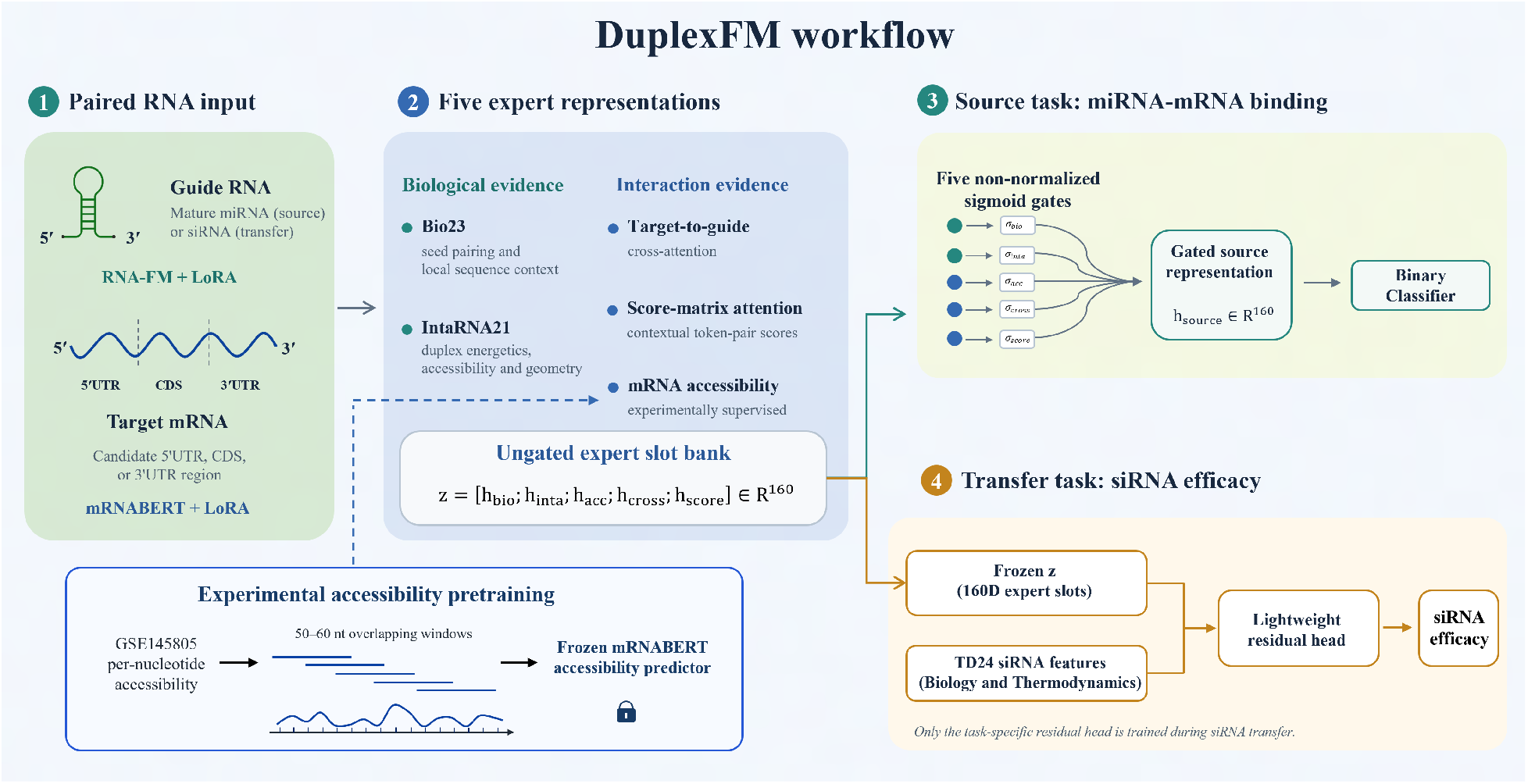
Overview of the DuplexFM workflow. LoRA-adapted RNA foundation models encode paired guide and target sequences, while five experts capture complementary biological, biophysical, accessibility, and sequence-interaction evidence. The resulting expert slots form *z*, which is adaptively gated for miRNA–mRNA binding prediction. For siRNA transfer, *z* is frozen and combined with TD24 features through a lightweight residual head to predict efficacy.

On the updated, bias-corrected miRBench v7 benchmark [13], the three-seed DuplexFM ensemble achieved the highest macro APS of 0.876 and outperformed miRBenchCNN-Manakov, the strongest baseline by APS, on all four test partitions. The learned gate profiles placed the greatest emphasis on Bio23 and IntaRNA21 and showed increased reliance on accessibility in the family-held-out setting. When transferred to six siRNA evaluation settings, an ensemble of three independently initialized transfer heads achieved the highest macro Pearson correlation, Spearman correlation, AUPRC, and F1 among the evaluated approaches, reaching 0.577, 0.597, 0.799, and 0.738, respectively. These findings indicate that miRNA-derived interaction representations capture transferable regulatory signals and provide a parameter-efficient means of augmenting limited siRNA-efficacy data.

The main contributions of this work are as follows:

- We introduce a biologically grounded, mechanism-prioritized multi-expert framework that integrates RNA foundation-model representations with complementary biological, thermodynamic, and accessibility evidence. All five expert representations are independently modulated by sample-specific sigmoid gates, with Bio23 and IntaRNA21 initialized as higher-prior core experts. On miRBench v7, the three-seed soft-voting ensemble achieved the highest macro APS of 0.877 and outperformed miRBenchCNN-Manakov, the strongest baseline by macro APS, on all four test partitions.
- We develop an experimentally supervised mRNA-accessibility expert using nucleotide-resolution icSHAPE measurements. Rather than representing accessibility by a fragment-wide average, the frozen predictor aligns its nucleotide-level predictions with Bio23-defined seed sites, IntaRNA-predicted pairing positions, and local target contexts. This provides a direct estimate of whether sequence- or energy-supported candidate binding regions are structurally accessible.
- We establish that interaction representations learned from large-scale miRNA–mRNA data are transferable to siRNA efficacy prediction. The frozen DuplexFM representation is adapted using a TD24-anchored residual head with only 6,387 trainable parameters. It consistently improves correlation and ranking performance over the matched TD24-only control across all six siRNA evaluation settings. An ensemble of independently initialized transfer heads further achieved the highest six-setting macro Pearson correlation, Spearman correlation, AUPRC, and F1 among the evaluated approaches.

## 2 Methods

### 2.1 Data Collection and Preprocessing

The data used in our work are presented in Table 1.To obtain site-level miRNA-mRNA interactions, we downloaded miRBench data [13]. miRBench is a benchmark dataset and evaluation framework for miRNA binding site prediction. It proposes a negative sample construction strategy that preserves the miRNA family distribution between positive and negative classes while sampling negatives from target-site clusters that do not overlap with the positive clusters of the same family, thereby reducing miRNA frequency class bias. miRBench offers processed chimeric eCLIP data from Manakov et al. [12] and AGO2 eCLIP data from Klimentová et al. [40], and AGO CLASH data from Hejret et al. [41]. Chimeric eCLIP is an AGO2 eCLIP-derived method that adds an RNA ligation step to capture miRNA–target chimeric reads, allowing direct identification of miRNA–RNA interaction pairs [12]. AGO2 eCLIP identifies AGO2-bound RNA regions transcriptome-wide, but does not by itself directly resolve the paired miRNA for each target site [42]. AGO2-CLASH directly maps miRNA–target duplexes by ligating AGO2-associated guide and target RNAs prior to sequencing [43].

**Table 1:**
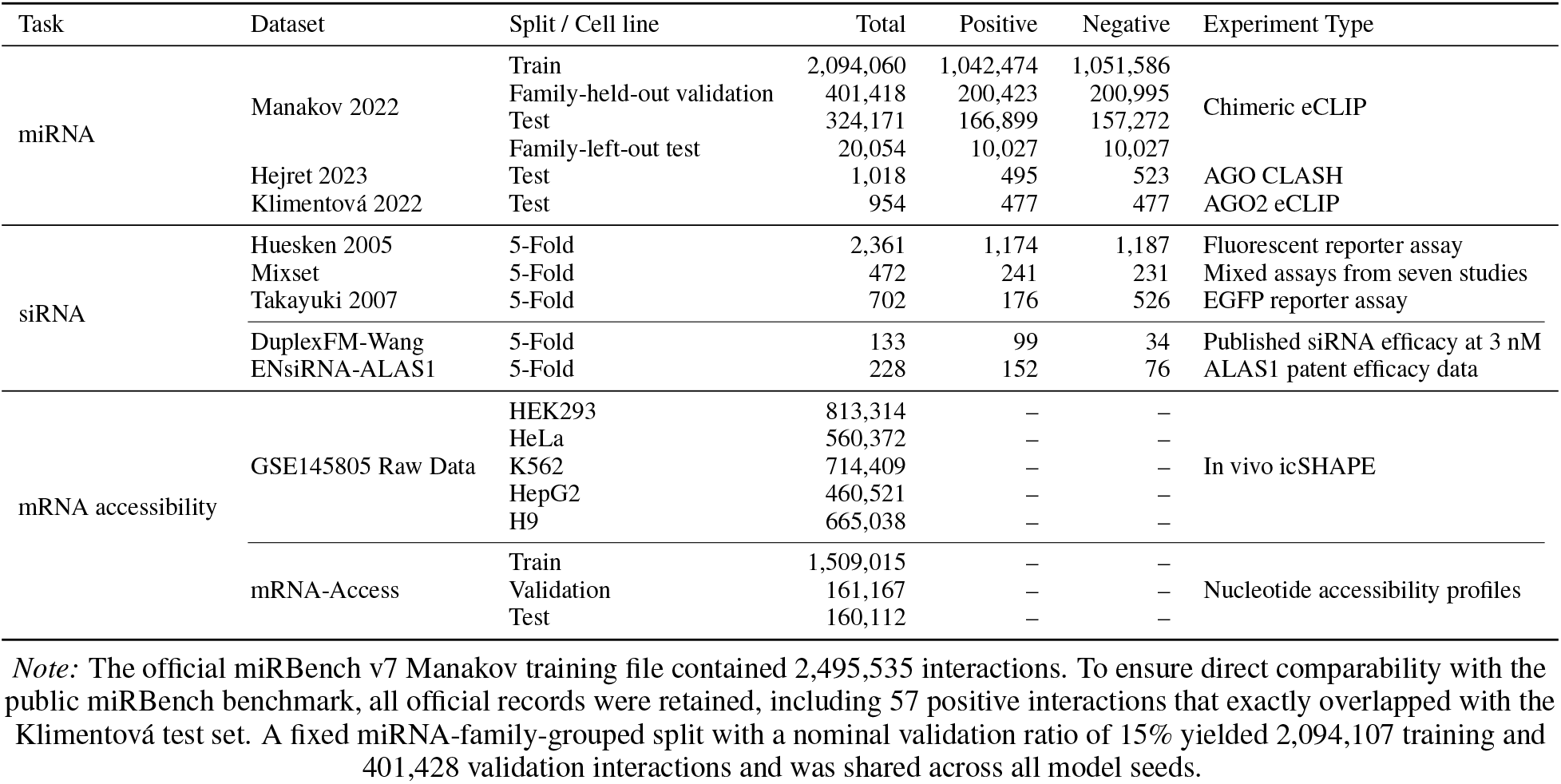
Dataset overview of this study.

To obtain siRNA efficacy data, we reused the datasets compiled by OligoFormer. OligoFormer employed datasets originally derived from nine sources, including Huesken et al. (2005), Katoh and Suzuki (2007), Amarzguioui et al. (2003), Harborth et al. (2003), Hsieh et al. (2004), Reynolds et al. (2004), Vickers et al. (2003), and Ui-Tei et al. (2004) [31, 44, 45, 46, 47, 27, 48, 28], comprising a total of 3,714 siRNAs and 75 mRNAs. For all datasets, the inhibition efficacy (activity) of siRNAs was normalized to a scale of 0% to 100%. A threshold corresponding to 70% of the original maximum inhibition was then applied to classify siRNAs into positive and negative categories [14].

To provide additional external evaluation sets, we additionally reconstructed an siRNA efficacy dataset from published supplementary materials. Because siRNA guides bind their mRNA targets antiparallelly, the transcript-derived target core, when written 5’ to 3’, was required to match the reverse complement of the guide after T/U normalization. The verified target site was then extended with transcript-derived flanking sequences to construct the model input window. The first dataset, hereafter referred to as DuplexFM-Wang, was reconstructed from Supplementary Table 3 of Wang et al. [49]. Repeated measurements of identical duplexes were averaged, and two lower-confidence transcript-context surrogates were excluded, resulting in 133 unique siRNA duplexes targeting 14 human genes. Their target contexts were recovered by mapping the reported 19-nt target cores to current or historically matched RefSeq transcripts, and siRNA efficacy measured at a concentration of 3 nM was used. For the second dataset, we reused the 232-entry ALAS1-targeting patent dataset released with ENsiRNA, hereafter referred to as ENsiRNA-ALAS1 [50]. Four entries whose centered 57-nt target windows contained ambiguous N bases were removed, leaving 228 siRNAs for evaluation. Continuous efficacy values were retained for correlation analyses, whereas efficacy thresholds of 0.70 for DuplexFM-Wang and 0.7 for ENsiRNA-ALAS1 were used for binary classification.

To construct the mRNA-accessibility dataset, we obtained nucleotide-resolution in vivo icSHAPE profiles from GSE145805 [51] and used icSHAPE reactivity as an experimental proxy for local RNA accessibility [52]. We retained five human cell lines, HEK293, HeLa, K562, HepG2, and H9, and excluded mouse data. Versioned Ensembl transcript identifiers were matched exactly to a GRCh38 reference, retaining 338,731 of 339,213 records whose declared length, reference length, and number of measurements agreed. Transcripts were partitioned into deterministic, non-overlapping 50–60-nt canonical fragments using seed 42. Terminal fragments shorter than 50 nt and fragments containing ambiguous bases, missing values, or non-finite measurements were discarded. Measurements for identical sequences across transcripts and cell lines were aggregated by their position-wise median, yielding 1,075,597 canonical sequences. Transcripts sharing an identical canonical fragment were assigned to the same data partition, producing 754,318 training, 161,167 validation, and 160,112 test sequences. To reduce boundary effects, a second segmentation offset by 27 nt was generated only from training transcripts; shifted sequences matching any canonical sequence were excluded. This added 754,697 training sequences, giving 1,509,015 training sequence-view records in total, while validation and test sets remained canonical-only. The resulting nucleotide-level profiles were used as continuous targets for LoRA adaptation of mRNABERT.

### 2.2 Model Construction

DuplexFM was implemented as a two-stage framework (Fig. 1). In the source stage, five complementary experts captured biological pairing, interaction energetics, target accessibility, and foundation-model-derived guide–target recognition. Their representations were adaptively fused to predict site-level miRNA–mRNA interactions. In the transfer stage, the five trained 32-dimensional expert slots were frozen and concatenated into a 160-dimensional representation. A lightweight TD24-anchored residual head then used this representation to predict continuous siRNA efficacy.

#### 2.2.1 Sequence representations with RNA foundation models

For each sample, the mature miRNA was treated as the guide and its local transcript fragment as the target. During siRNA transfer, the miRNA was replaced by the siRNA guide. We retained target fragments from all annotated transcript regions represented in miRBench v7. RNA-FM [35] encoded the guide, whereas mRNABERT [39] encoded the target. Both models operated at single-nucleotide resolution. The maximum tokenized lengths were 32 tokens for the guide and 64 tokens for the target, including boundary special tokens; padding tokens were masked.

The cross-attention and score-matrix experts used separate RNA-FM/mRNABERT encoder pairs. Both pairs were initialized from the same public pretrained checkpoints ^1^, but did not share task-specific parameters. The pretrained backbone weights were frozen and adapted using branch-specific LoRA parameters. For both backbones, LoRA used rank *r* = 16, scaling factor *α* = 32, and dropout 0.05. It was applied to the fused query–key–value projection of mRNABERT and the separate query, key, and value projections of RNA-FM. The independently trained mRNA-accessibility predictor also used mRNABERT but remained frozen during miRNA and siRNA modeling, as described below.

#### 2.2.2 Five-expert representation bank

The representation bank comprised Bio23, IntaRNA21, mRNA accessibility, mRNA-to-miRNA/siRNA cross-attention, and score-matrix attention. Bio23 and IntaRNA21 were treated as always-on *core* experts because they directly describe pairing rules and interaction energetics. The accessibility, cross-attention, and score-matrix experts were treated as sample-dependent *plugins*. Their raw feature dimensions were 23, 21, 10, 256, and 64, respectively.

##### Bio23 biological expert

Bio23 encoded 23 biologically motivated features describing canonical seed-site abundance and quality, local AU context, position-resolved Watson–Crick and G:U pairing, noncanonical and extended pairing patterns, target A1, and canonical/offset site density. These features summarized both established seed rules and additional pairing configurations, including supplementary, compensatory, centered, and single-gap interactions. The same four canonical-site definitions were also used to identify the seed-conditioned nucleotide-level profiles in the accessibility expert.

##### IntaRNA21 biophysical expert

For each guide-target pair, IntaRNA v3.4.1 [53] was applied to the mRNA target fragment and the full guide sequence in heuristic pairwise mode, without imposing a seed constraint. We retained at most one interaction: the top-ranked prediction with a reported total energy of *E* ≤ 0. The interaction was represented by 21 interpretable features describing binding energetics, the energetic costs of exposing the target and guide, and the extent, composition, and continuity of duplex pairing. A binary indicator distinguished pairs for which no valid interaction was returned. The target-side coordinates of the retained base pairs were also used to align the frozen nucleotide-level accessibility predictions with the proposed duplex. Finally, *x*_inta_ ∈ ℝ^21^ was projected into a 32-dimensional expert slot and modulated by an independently learned sample-specific sigmoid gate.

##### mRNA accessibility plugin

Sequence complementarity alone is often insufficient to ensure small-RNA binding, because a candidate target site may be sequestered by local mRNA structure and thus remain inaccessible to the guide RNA [54, 55]. Motivated by this, we developed *mRNA-Access*, an experimentally supervised predictor that estimates nucleotide-resolution target accessibility from icSHAPE data processed from the human GSE145805 dataset. For identical RNA sequences measured across the HEK293, HeLa, K562, HepG2, and H9 cell lines, the per-nucleotide values were aggregated by taking the median. To construct the training set, each icSHAPE-annotated mRNA was first partitioned into consecutive, variable-length fragments of 50–60 nt. In addition, a second set of fragments of the same 50–60 nt length was generated by sliding a window with a 27-nt stride along each mRNA, providing shifted, overlapping views. The predictor was then trained jointly on the original consecutively partitioned fragments and these shifted fragments. This strategy can reduce sensitivity to artificial window boundaries.

mRNA-Access used mRNABERT with rank-16 LoRA adaptation [56]. In addition to the nucleotide representations, we retained the mRNABERT CLS and SEP states because these boundary tokens can carry fragment-level sequence context. Their representations were used only to generate feature-wise scaling and shifting factors. CLS and SEP were not assigned accessibility labels and did not enter the nucleotide-level predictions. The conditioned nucleotide representations were processed by a position-shared direct prediction branch and a lightweight two-layer residual Transformer, producing an accessibility score *a*_*i*_ ∈ [0, 1] for each nucleotide.

Rather than reducing accessibility to a single fragment-wide average, we summarized the predicted profile over regions directly relevant to small-RNA binding:

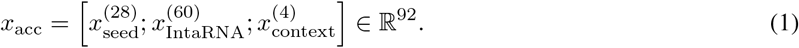

The 28 seed-conditioned values retained the single-nucleotide-level accessibility profiles of the most accessible canonical 6mer, 7mer-A1, 7mer-m8, and 8mer sites. The 60 IntaRNA-conditioned values represented accessibility at target nucleotides participating in the top-ranked IntaRNA duplex, using a centered coordinate system that supports target fragments of different lengths. Unpaired, padded, or unavailable positions were set to zero. The remaining four features summarized accessibility over the central 25-nt region, its upstream and downstream flanks, and the complete target fragment. This design allowed the accessibility expert to assess whether sequence- or energy-supported candidate pairing regions were locally open while retaining broader structural context.

During miRNA source-task training and siRNA transfer, only the adapter Linear(92, 32)-GELU-Dropout-LayerNorm and the downstream task-specific components were trainable. The mRNA-Access checkpoint remained frozen and was used to generate static accessibility features. It was therefore independent of the separately initialized mRNABERT encoders used by the cross-attention and score-matrix experts.

##### mRNA-to-small-RNA cross-attention plugin

This plugin modeled guide-target recognition asymmetrically by asking which parts of the miRNA/siRNA guide were informative for each position in the candidate mRNA target. The mRNABERT target representations and RNA-FM guide representations were first projected into a shared 256-dimensional space. The mRNA tokens then served as queries, whereas the guide tokens served as keys and values. The resulting guide-derived signal was added back to the original target representation:

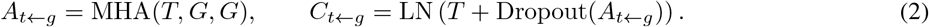

This residual update allowed each target position to retain its original mRNA context while incorporating information from the guide. We summarized the variable-length representation using two complementary readouts: learned attention pooling emphasized the most informative target positions, whereas masked mean pooling retained the overall interaction context. Their concatenation was compressed to a 256-dimensional vector and subsequently mapped to the 32-dimensional cross-attention expert slot. The slot was then modulated by its sample-specific sigmoid gate during expert fusion.

##### Score-matrix attention plugin

A separate, parameter-unshared RNA-FM/mRNABERT LoRA encoder pair projected the guide and target tokens into a shared 256-dimensional interaction space. Their contextual compatibility was represented by

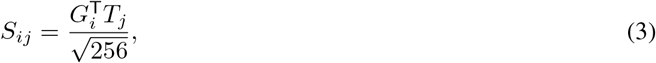

where each matrix cell describes the compatibility between one guide token and one target token. Only nucleotide–nucleotide cells were retained as candidate interactions. Boundary special tokens remained available to the pretrained encoders and provided global guide–target context, but were excluded from the candidate-pair mask.

Each scalar compatibility score was transformed nonlinearly into a 64-dimensional representation and augmented with separate guide- and target-position embeddings. These pair representations were further modulated by the global sequence context, allowing similar compatibility scores to be interpreted differently depending on their positions and the overall guide–target pair. Learned masked attention pooling over all valid cells produced *x*_score_ ∈ ℝ^64^, which was subsequently mapped to the 32-dimensional score-matrix expert slot. No explicit Watson–Crick, G:U, or seed-pairing rules were imposed in this plugin.

#### 2.2.3 Adaptive expert fusion and miRNA-target prediction

The five experts capture complementary aspects of small-RNA targeting and differ substantially in scale and dimensionality. Each expert representation was therefore mapped to an independent 32-dimensional slot. A shared router examined all five slots jointly and assigned each expert a sample-specific sigmoid gate. Unlike softmax routing, these gates were not constrained to sum to one, because seed pairing, favorable interaction energy, target accessibility, and sequence-context evidence may all be informative for the same interaction. Bio23 and IntaRNA21 were designated as core experts and initialized with higher gate values to reflect their direct biological relevance. Nevertheless, all five gates remained learnable. The gated slots were concatenated into a 160-dimensional representation and passed to a binary classifier to estimate the probability of miRNA–mRNA interaction.

#### 2.2.4 Source-task optimization

DuplexFM was trained jointly on the all-region Manakov 2022 partition of miRBench v7. The branch-specific LoRA parameters and interaction layers of the cross-attention and score-matrix experts were optimized together with the five expert adapters, router, and classifier. In contrast, the experimentally trained mRNA-Access predictor remained frozen, allowing its accessibility estimates to serve as an external structural prior. A weak auxiliary classification head was attached to each expert during training to preserve interaction-relevant information within every slot. These auxiliary heads were discarded after training. Checkpoints were selected exclusively by validation APS and subsequently evaluated on the four miRBench test sets using strict FP32 inference. Three independently initialized models were trained with seeds 42-44, and their predicted probabilities were averaged for the soft-voting ensemble.

#### 2.2.5 Frozen transfer to siRNA efficacy prediction

To investigate whether interaction knowledge learned from miRNA data could support siRNA efficacy prediction, we froze the five adapted source-expert slots and discarded the miRNA-specific router, auxiliary heads, and classifier. The ungated slots were concatenated into a 160-dimensional representation *z*, thereby retaining the complete source evidence without imposing routing decisions learned specifically for miRNA binding. TD24 descriptors *t* were used to establish an siRNA-specific efficacy baseline, and the frozen DuplexFM representation provided a lightweight residual correction:

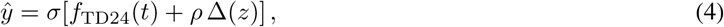

where *f*_TD24_ is the task-specific anchor, Δ(*z*) reads the frozen source representation, and *ρ* is a learned bounded global coefficient. Only the TD24 anchor, residual head, and *ρ* were optimized on the siRNA data. Training used continuous efficacy regression with a penalty on the magnitude of the source-derived correction:

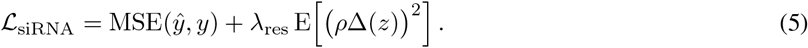

No binary classification loss was used. This formulation directly tests whether miRNA-derived interaction representations provide information complementary to established siRNA-specific descriptors.

## 3 Results

### 3.1 LoRA-adapted mRNABERT captures nucleotide-resolved mRNA accessibility

For sequences measured in multiple human cell lines in GSE145805, the nucleotide-level target was defined as the median icSHAPE value across cell lines. The transcript-component-disjoint split comprised 754,318 canonical training fragments, 161,167 validation fragments, and 160,112 test fragments. To reduce sensitivity to artificial fragment boundaries, we generated an additional 754,697 training-only views by repeating the 50-60-nt partition from a 27-nt offset. This yielded 1,509,015 training views in total, whereas validation and testing used only the original canonical fragments. The resulting mRNA-Access model combined LoRA-adapted mRNABERT with CLS/SEP-conditioned feature modulation and a lightweight two-layer residual Transformer to predict one accessibility value per nucleotide.

On the held-out test set, mRNA-Access achieved a Pearson correlation of 0.627, a Spearman correlation of 0.624, and a mean absolute error of 0.160 across 8,800,970 valid nucleotide positions (Fig. 2a). Among the 160,029 test fragments with finite correlations, the mean within-sequence Pearson correlation was 0.645, and the median was 0.674. These results indicate that the model captured relative accessibility variation along individual mRNA fragments rather than relying only on differences in mean accessibility between sequences.

**Figure 2:**
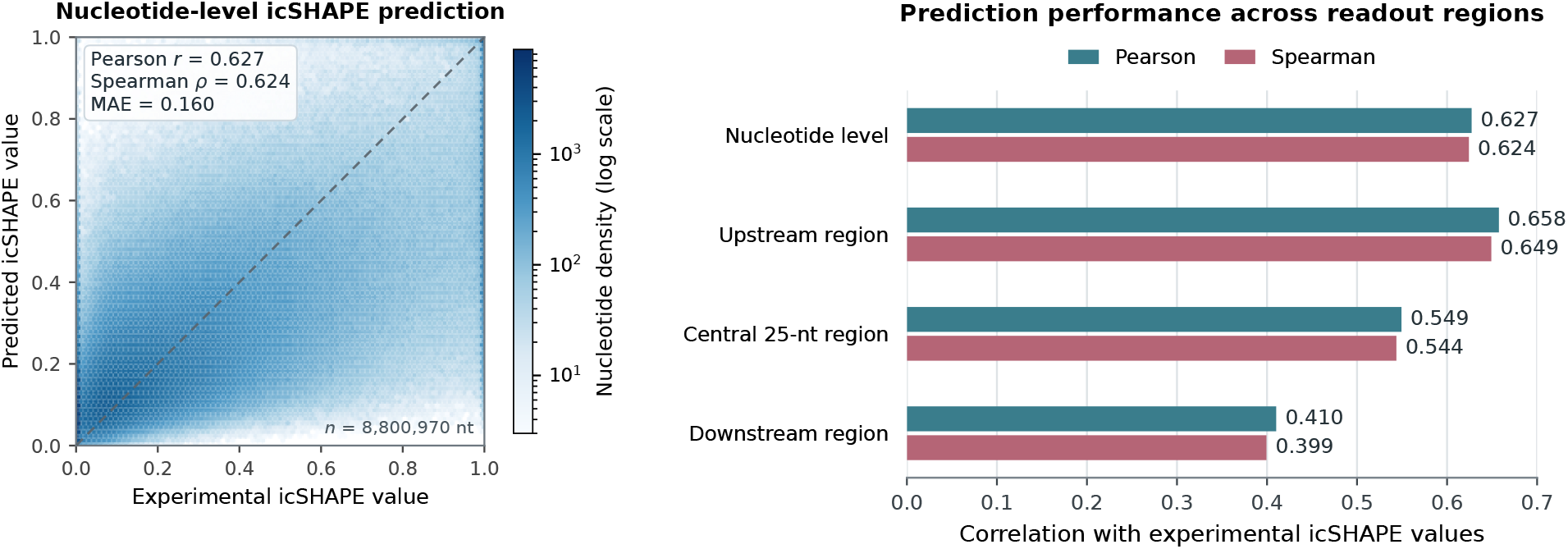
Performance of the LoRA-adapted mRNABERT accessibility model on the transcript-component-disjoint human GSE145805 test set. Left, Hexagonal density of experimental and predicted nucleotide-level icSHAPE values. A deterministic sample of 1.5 million positions is displayed for visualization, whereas the reported statistics were calculated over all 8,800,970 valid nucleotide positions. Right, Pearson and Spearman correlations at the nucleotide level and after averaging experimental and predicted icSHAPE values within the upstream, central 25-nt, and downstream regions.

**Figure 3:**
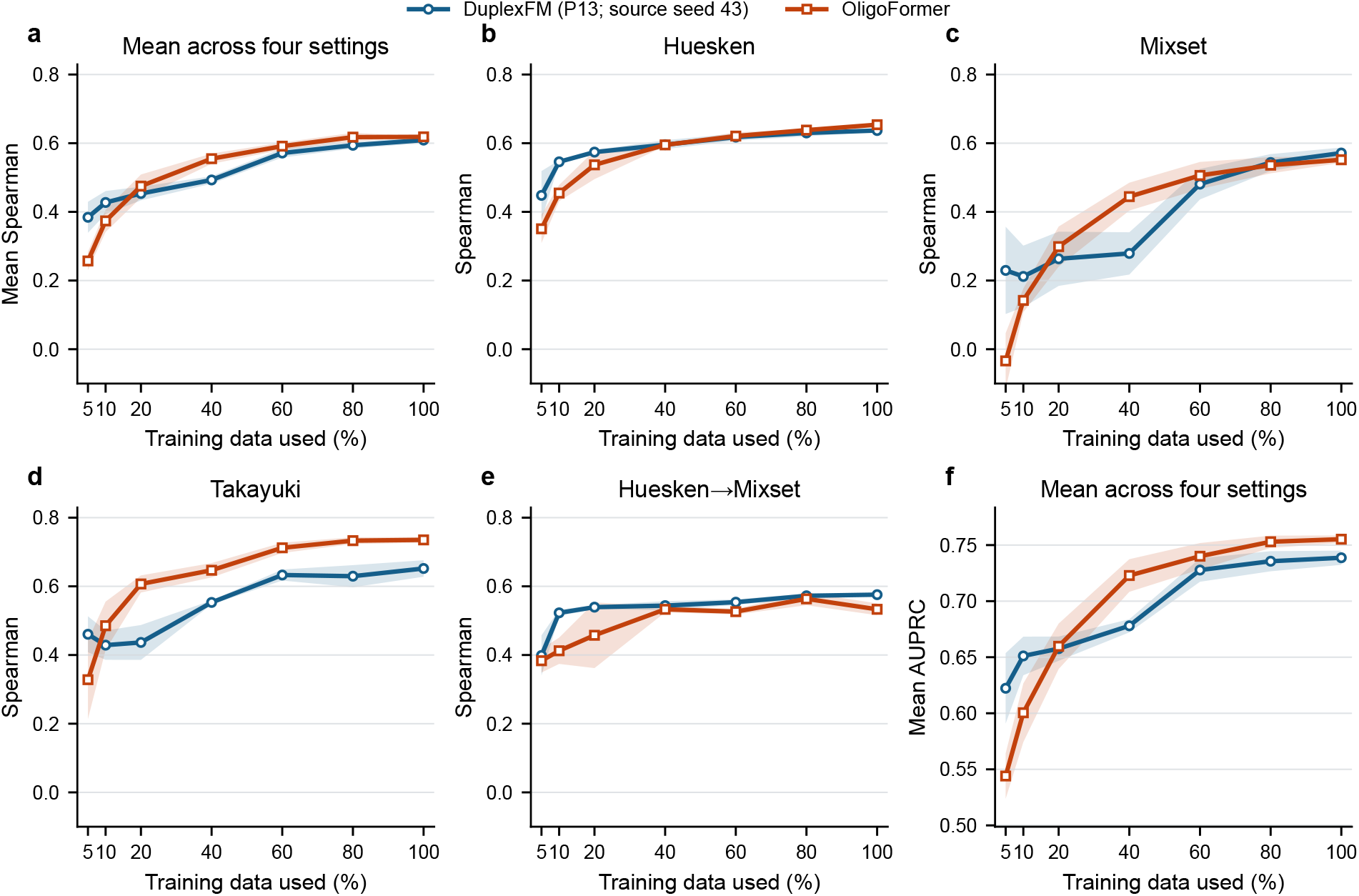
Effect of siRNA training-set size on DuplexFM transfer and OligoFormer performance. Training fractions ranged from 5% to 100%, with the outer test partitions held fixed across all fractions. Panels show (a) the unweighted mean Spearman correlation across the four evaluation settings; Spearman correlation on (b) Huesken, (c) Mixset, (d) Takayuki, and (e) Huesken→Mixset; and (f) the mean AUPRC across the four settings. For DuplexFM, the miRNA-trained expert slots from source seed 43 were frozen, and only the TD24-anchored P13 full-slot residual head was trained. DuplexFM curves show the mean across transfer-head initialization seeds 42–44, whereas OligoFormer curves show the mean across independently initialized model seeds 42–44; shaded regions indicate ± one standard deviation. At the aggregate level, DuplexFM achieved its largest advantage at the 5% and 10% training fractions, whereas OligoFormer matched or exceeded its performance from 20% onward.

Regional aggregation revealed substantial positional variation in predictive performance (Fig. 2b). After accessibility was averaged within each region of every fragment, the upstream region achieved a Pearson correlation of 0.658. The corresponding correlations were 0.549 for the central 25-nt candidate-binding region and 0.410 for the downstream region. Thus, the model retained informative accessibility signals within the candidate-binding region, although prediction accuracy remained dependent on position within the target fragment.

Because accessibility alone does not establish guide-target recognition, we incorporated mRNA-Access into DuplexFM as a frozen source of complementary evidence. Its nucleotide-level predictions were organized into a 92-dimensional representation comprising 28 values for the most accessible canonical 6mer, 7mer-A1, 7mer-m8, and 8mer seed sites. 60 centered values for target nucleotides paired in the top-ranked IntaRNA duplex, and four regional summaries covering the central 25-nt region, its upstream and downstream flanks, and the complete target fragment. All mRNA-Access parameters remained frozen during subsequent DuplexFM training and transfer. Only the adapter mapping the 92-dimensional representation into its 32-dimensional expert slot and the downstream task-specific components were optimized.

### 3.2 Benchmarking in miRBench

We compared DuplexFM with the predictors reported in the current miRBench version 7 benchmark [13]. Public baseline scores were obtained from the four official miRBench test partitions. For direct comparison under the official benchmark protocol, DuplexFM used the unmodified Manakov training partition. All three training runs used the same fixed, family-grouped validation split and differed only in their training seeds. The selected checkpoints were evaluated using strict FP32 inference.

To summarize performance independently of test-set size, we calculated macro APS as the unweighted average of the four test-set APS values. Across seeds 42, 43, and 44, DuplexFM achieved a macro APS of 0.873 ± 0.002, exceeding the 0.868 obtained by the strongest public baseline, miRBenchCNN_Manakov (Table 2). The mean single-model performance was highest on Klimentova2022 and was within 0.004, 0.001, and 0.001 of the strongest public results on Hejret2023, Manakov2022 test, and Manakov2022 left-out, respectively. Averaging the prediction probabilities of the three DuplexFM models further increased macro APS to 0.876. This soft-voting ensemble achieved the highest APS on all four test partitions, reaching 0.883, 0.889, 0.872, and 0.861, respectively.

**Table 2:** Performance comparison on miRBench version 7 using average precision score (APS). Macro APS is the unweighted average across the four test sets. Public baseline results were obtained from miRBench. DuplexFM results are reported as the mean ± sample standard deviation across training seeds 42, 43, and 44, using strict FP32 inference. The three-seed soft-voting ensemble averages the prediction probabilities of these three independently trained models and therefore does not report a standard deviation. The highest value in each column is shown in bold.

| Model | Macro APS | Klimentova2022 test | Hejret2023 test | Manakov2022 test | Manakov2022 leftout |
| --- | --- | --- | --- | --- | --- |
| Random | 0.520 | 0.534 | 0.526 | 0.515 | 0.505 |
| CnnMirTarget_Zheng2020 | 0.515 | 0.515 | 0.511 | 0.526 | 0.508 |
| TargetNet_Min2021 | 0.568 | 0.547 | 0.553 | 0.576 | 0.596 |
| RNACofold | 0.683 | 0.685 | 0.742 | 0.638 | 0.669 |
| InteractionAwareModel_Yang2024 | 0.700 | 0.702 | 0.741 | 0.712 | 0.644 |
| miRNA_CNN_Hejret2023 | 0.755 | 0.758 | 0.795 | 0.731 | 0.738 |
| miRBind_Klimentova2022 | 0.763 | 0.777 | 0.799 | 0.732 | 0.745 |
| TargetScanCnn_McGeary2019 | 0.784 | 0.787 | 0.742 | 0.800 | 0.807 |
| miRBenchCNN_HejretCorrected | 0.828 | 0.801 | 0.888 | 0.807 | 0.818 |
| miRBenchCNN_Manakov | 0.868 | 0.873 | 0.869 | 0.870 | 0.860 |
| DuplexFM | 0.873 ± 0.002 | 0.879 ± 0.005 | 0.884 ± 0.003 | 0.869 ± 0.002 | 0.859 ± 0.001 |
| DuplexFM (three-seed soft-voting) | **0.876** | **0.883** | **0.889** | **0.872** | **0.861** |

### 3.3 DuplexFM exhibits dataset-dependent expert routing

We next examined how DuplexFM routed information from the five experts across the miRBench test partitions. In the final architecture, all five expert slots were modulated by sample-specific sigmoid gates, including the two experts designated as biological cores. The router jointly considered the five ungated expert representations and produced five non-normalized coefficients. For each model seed, we first averaged each gate over all samples within a test partition. Table 3 reports the mean and standard deviation of these dataset-level values across seeds 42, 43, and 44.

**Table 3:** Dataset-level expert-gate distributions in DuplexFM. For each training seed, gate values were first averaged over all samples in the indicated miRBench test partition. Values are the mean ± sample standard deviation of these averages across seeds 42, 43, and 44. Each gate is a separately sigmoid-transformed coefficient in (0, 1); the five coefficients are not normalized and do not sum to one.

| Dataset | Bio23 | IntaRNA21 | Accessibility | Cross-attention | Score-matrix |
| --- | --- | --- | --- | --- | --- |
| Klimentova2022 test | 0.657 ± 0.061 | 0.478 ± 0.055 | 0.176 ± 0.022 | 0.398 ± 0.040 | 0.464 ± 0.077 |
| Hejret2023 test | 0.720 ± 0.054 | 0.523 ± 0.042 | 0.170 ± 0.025 | 0.359 ± 0.031 | 0.416 ± 0.077 |
| Manakov2022 test | 0.705 ± 0.057 | 0.480 ± 0.064 | 0.165 ± 0.026 | 0.393 ± 0.051 | 0.449 ± 0.074 |
| Manakov2022 left-out | 0.772 ± 0.050 | 0.526 ± 0.058 | 0.196 ± 0.021 | 0.334 ± 0.029 | 0.372 ± 0.049 |

Bio23 consistently received the highest mean gate, ranging from 0.657 on Klimentova2022 to 0.772 on the family-held-out Manakov2022 partition. IntaRNA21 remained moderately activated across datasets, with mean gates between 0.478 and 0.526. Among the three plugin experts, score-matrix attention received the highest mean gate on every partition (0.372–0.464), followed by target-to-guide cross-attention (0.334–0.398). The mean accessibility gate was lower (0.165–0.196), indicating more selective use of this signal. Nevertheless, its sample-level distributions were right-skewed: after averaging the seed-specific percentiles, its 95th percentile ranged from 0.355 to 0.445 across the four datasets. Thus, accessibility was retained more strongly for a subset of guide-target pairs rather than being applied uniformly.

The routing pattern also changed under dataset shift. On the family-held-out Manakov2022 partition, Bio23 and accessibility reached their highest mean gates, whereas cross-attention and score-matrix attention reached their lowest. In contrast, Klimentova2022 produced the highest mean cross-attention and score-matrix gates and the lowest Bio23 gate. These patterns suggest that DuplexFM shifted its routing toward biologically grounded and accessibility-related evidence when generalizing to unseen miRNA families, while relying more strongly on learned interaction representations in the other test settings.

Because the five gates were not constrained to sum to one, they should not be interpreted as proportions of model attention. Their absolute values were also influenced by the different initialization probabilities used for core and plugin experts and by the scale and direction of the corresponding 32-dimensional expert representations. The gate distributions therefore describe sample-dependent routing behavior rather than the causal importance of individual experts.

### 3.4 Transferability to the siRNA Task

We next investigated whether interaction representations learned from miRNA–mRNA binding could support siRNA efficacy prediction. The final seed-43 DuplexFM checkpoint was used as the source model, and all source parameters remained frozen. The five 32-dimensional expert slots were concatenated into a 160-dimensional representation, whereas the source-task router and classifier were discarded. This frozen representation was supplied to a lightweight TD24-anchored residual head, which learned a correction to the prediction generated from the siRNA-specific TD24 features. The transfer head was optimized using continuous efficacy regression with residual regularization and without a binary auxiliary loss. Only 6,387 parameters were trainable in each DuplexFM transfer head, compared with 881 parameters in the TD24-only control.

Evaluation comprised five-fold experiments on Huesken, Mixset, and Takayuki, as well as an inter-dataset experiment in which models were trained on Huesken and evaluated on Mixset. DuplexFM-Wang and ENsiRNA-ALAS1 were evaluated by grouped five-fold out-of-fold prediction. The frozen source checkpoint was fixed throughout, whereas the task-specific heads were independently initialized using seeds 42, 43, and 44. The TD24-only and DuplexFM transfer models used identical outer folds, inner validation partitions, initialization seeds, feature normalization, and checkpoint selection rules. They therefore differed only in whether the frozen DuplexFM representation was available to the prediction head.

The frozen DuplexFM representation consistently provided information beyond the biological and thermodynamic TD24 descriptors. Averaged across the three head-initialization seeds, DuplexFM transfer improved Pearson correlation, Spearman correlation, AUPRC, and F1 over the TD24-only control in all six evaluation settings (Table 4). Across the six settings, macro Pearson correlation increased from 0.482 to 0.535, Spearman correlation from 0.504 to 0.552, AUPRC from 0.757 to 0.782, and F1 from 0.708 to 0.723. The absolute improvements ranged from 0.005 to 0.097 for Pearson correlation and from 0.014 to 0.100 for Spearman correlation. The largest gains were observed on ENsiRNA-ALAS1, where Pearson and Spearman correlations increased by 0.097 and 0.100, respectively.

**Table 4:** Transfer performance on four OligoFormer evaluation settings and two external siRNA datasets. Non-ensemble results are means over three random seeds and are rounded to three decimal places. For DuplexFM transfer, the source checkpoint was fixed at seed 43, whereas the prediction-head initialization seeds were 42–44. TD24 only and DuplexFM transfer used identical data partitions, normalization, initialization seeds, and model-selection rules. Ensemble rows were obtained by averaging the three continuous efficacy predictions within each evaluation split before calculating the out-of-fold metrics. The macro average is the unweighted mean across all six settings. The highest value in each column within each setting is shown in bold.

| Dataset | Method | Pearson | Spearman | AUPRC | F1 |
| --- | --- | --- | --- | --- | --- |
| Huesken | miRBenchCNN-Manakov (retrain) | 0.018 | 0.021 | 0.505 | 0.653 |
|  | TD24 only | 0.609 | 0.616 | 0.801 | 0.744 |
|  | OligoFormer | **0.647** | **0.654** | 0.810 | **0.758** |
|  | DuplexFM transfer | 0.626 | 0.634 | 0.810 | 0.754 |
|  | DuplexFM transfer (three-seed ensemble) | 0.633 | 0.640 | **0.815** | 0.755 |
| Mixset | miRBenchCNN-Manakov (retrain) | 0.022 | 0.017 | 0.513 | 0.676 |
|  | TD24 only | 0.479 | 0.502 | 0.748 | 0.715 |
|  | OligoFormer | 0.538 | 0.552 | 0.763 | 0.716 |
|  | DuplexFM transfer | 0.556 | 0.570 | 0.782 | **0.720** |
|  | DuplexFM transfer (three-seed ensemble) | **0.580** | **0.595** | **0.790** | 0.713 |
| Takayuki | miRBenchCNN-Manakov (retrain) | 0.022 | 0.039 | 0.256 | 0.401 |
|  | TD24 only | 0.599 | 0.607 | 0.516 | 0.514 |
|  | OligoFormer | **0.739** | **0.735** | **0.681** | **0.544** |
|  | DuplexFM transfer | 0.637 | 0.642 | 0.552 | 0.524 |
|  | DuplexFM transfer (three-seed ensemble) | 0.671 | 0.673 | 0.591 | 0.531 |
| Huesken→Mixset | miRBenchCNN-Manakov (retrain) | 0.014 | 0.029 | 0.499 | 0.676 |
|  | TD24 only | 0.556 | 0.555 | 0.781 | 0.741 |
|  | OligoFormer | 0.536 | 0.533 | 0.768 | **0.748** |
|  | DuplexFM transfer | 0.562 | 0.569 | 0.789 | 0.742 |
|  | DuplexFM transfer (three-seed ensemble) | **0.567** | **0.574** | **0.792** | 0.747 |
| DuplexFM-Wang | miRBenchCNN-Manakov (retrain)<sup>†</sup> | −0.040 | 0.001 | 0.739 | 0.853 |
|  | TD24 only | 0.284 | 0.357 | 0.872 | 0.812 |
|  | OligoFormer | 0.181 | 0.243 | 0.798 | 0.843 |
|  | DuplexFM transfer | 0.362 | 0.416 | 0.899 | 0.834 |
|  | DuplexFM transfer (three-seed ensemble) | **0.459** | **0.518** | **0.919** | **0.870** |
| ENsiRNA-ALAS1 | miRBenchCNN-Manakov (retrain)<sup>†</sup> | −0.014 | −0.011 | 0.683 | 0.800 |
|  | TD24 only | 0.367 | 0.384 | 0.826 | 0.720 |
|  | OligoFormer | 0.480 | 0.488 | 0.840 | 0.806 |
|  | DuplexFM transfer | 0.464 | 0.484 | 0.859 | 0.762 |
|  | DuplexFM transfer (three-seed ensemble) | **0.549** | **0.580** | **0.885** | **0.812** |
| Macro average | miRBenchCNN-Manakov (retrain) | 0.004 | 0.016 | 0.533 | 0.677 |
|  | TD24 only | 0.482 | 0.504 | 0.757 | 0.708 |
|  | OligoFormer | 0.520 | 0.534 | 0.777 | 0.736 |
|  | DuplexFM transfer | 0.535 | 0.552 | 0.782 | 0.723 |
|  | DuplexFM transfer (three-seed ensemble) | **0.577** | **0.597** | **0.799** | **0.738** |

F1 used dataset-specific binary labels and a fixed prediction threshold of 0.5; correlation and AUPRC were treated as the primary transfer metrics. OligoFormer results were obtained from our local re-evaluation and differ from the published values [14] because we used the released processed data, prespecified splits, three-seed out-of-fold aggregation, and an audited sample-independent implementation. The complete reproduction and comparison code will be released with DuplexFM. <sup>†</sup>miRBenchCNN-Manakov classified every external sample as positive, yielding zero specificity; its F1 therefore largely reflects class prevalence (mean AUROC: 0.489 on DuplexFM-Wang and 0.499 on ENsiRNA-ALAS1).

We further evaluated whether prediction averaging could reduce sensitivity to head initialization. For each evaluation split, the continuous efficacy predictions from seeds 42–44 were averaged before calculating the metrics. To distinguish the contribution of the transferred representation from a generic ensemble effect, the same procedure was applied to the TD24-only control. The TD24-only ensemble achieved macro Pearson and Spearman correlations of 0.521 and 0.543, whereas the DuplexFM ensemble reached 0.577 and 0.597. Macro AUPRC increased from 0.772 to 0.799 and F1 from 0.721 to 0.738. The DuplexFM ensemble outperformed the matched TD24-only ensemble in all four metrics on every evaluation setting, showing that its improvement could not be explained by prediction averaging alone.

Among the non-ensemble models, DuplexFM transfer achieved higher six-setting macro Pearson correlation, Spearman correlation, and AUPRC than the task-specific OligoFormer model [14], whereas OligoFormer retained a higher macro F1. OligoFormer was stronger on Huesken and Takayuki, while DuplexFM transfer produced higher correlations on Mixset, Huesken-to-Mixset, and DuplexFM-Wang. The DuplexFM ensemble achieved the highest correlations in four of the six settings and the highest AUPRC in five. Because OligoFormer was summarized as the mean performance of independently trained models rather than as an ensemble, these ensemble comparisons were treated as secondary; the paired comparison between DuplexFM transfer and TD24 only constituted the primary test of transferability.

In contrast, the fully retrained miRBenchCNN-Manakov model produced correlations close to zero. Its high F1 values on the two external datasets resulted from predicting every sample as positive and therefore largely reflected class prevalence. Together, these results demonstrate that a frozen representation learned from miRNA–mRNA interactions contains complementary information for siRNA efficacy prediction and can be adapted using only a small task-specific prediction head.

## 4 Discussion

In this study, we developed DuplexFM to investigate whether knowledge learned from large-scale miRNA–mRNA interaction data can be transferred to siRNA efficacy prediction. DuplexFM achieved a macro APS of 0.876 across the four miRBench v7 test sets. These results indicate that RNA foundation-model representations can be effectively combined with pairing rules and interaction energetics. More importantly, the same representation could be transferred to siRNA prediction without retraining the source model, supporting the hypothesis that miRNA and siRNA targeting share learnable interaction patterns.

For siRNA transfer, DuplexFM was kept frozen and only a lightweight TD24-anchored residual head with 6,387 trainable parameters was optimized. This corresponds to approximately 0.44% of the trainable parameters in our local OligoFormer reproduction. The frozen DuplexFM representation consistently improved the matched TD24-only baseline, supporting the presence of interaction information that complements conventional siRNA descriptors.

DuplexFM was competitive with OligoFormer across the six evaluation settings, with clearer advantages on Mixset, inter-dataset transfer, and external validation, whereas OligoFormer remained stronger on Takayuki. Ensembling three transfer heads further improved robustness. These findings support parameter-efficient transfer of miRNA-derived interaction knowledge rather than universal superiority over a task-specific siRNA model. The parameter comparison concerns adaptation cost only, as DuplexFM still requires its frozen source encoders during feature generation.

Analysis of the learned expert gates provided further insight into how DuplexFM combined heterogeneous evidence. Across the test partitions, the model assigned the largest average gates to Bio23 and IntaRNA21, indicating that explicit pairing rules and thermodynamic and duplex-geometry features were the most consistently retained signals. Notably, on the family-held-out Manakov partition, the model placed greater emphasis on Bio23, IntaRNA21, and mRNA accessibility. This pattern is biologically reasonable: when encountering miRNA families absent from training, the model relied more strongly on mechanistically grounded evidence, such as whether a plausible duplex can form or whether the corresponding target region is accessible. Rather than solely on learned interaction patterns. Because gate values modulate expert representations and are not calibrated causal-importance scores, they should be interpreted as differences in model reliance rather than as a definitive ranking of biological relevance.

Several limitations remain. On the held-out test set, the accessibility predictor achieved a pooled nucleotide-level Pearson correlation of 0.627. When accessibility was averaged over the central 25-nt region of each fragment, the correlation between predicted and observed regional means was 0.549, indicating that substantial experimental variation remained unexplained. Moreover, using median icSHAPE profiles across five human cell lines removed cell-type-specific signals. Although shifted overlapping windows reduced boundary dependence, each prediction was still restricted to a 50-60-nt sequence and could not capture long-range interactions, RNA-binding proteins, modifications, or other cellular determinants of accessibility.

Prediction accuracy also varied by position. Regional-mean Pearson correlations were 0.658, 0.549, and 0.410 for the upstream, central, and downstream regions, respectively. This asymmetry may reflect residual boundary effects, positional or sequence-composition biases, and directional patterns learned during mRNABERT pretraining. In addition, icSHAPE reactivity is an in vivo proxy rather than a direct measure of thermodynamic unpaired probability. Finally, the selected accessibility architecture was evaluated using a single optimization seed. Future work should compare alternative encoders, longer-context models, and cell-type-aware targets under a controlled multi-seed protocol.

Overall, DuplexFM demonstrates that miRNA-mRNA interaction data can provide biologically relevant supervision for siRNA efficacy prediction. Rather than uniformly outperforming a dedicated siRNA model, its central contribution is to show that a frozen miRNA-derived representation provides information complementary to siRNA-specific descriptors and can be adapted using only a lightweight residual head. This offers a data-efficient strategy for leveraging abundant miRNA interaction measurements to support siRNA modeling, for which matched efficacy experiments remain costly and limited. More broadly, our findings suggest that miRNA and siRNA targeting share a transferable representation of small-RNA-mRNA recognition, providing a basis for unified models of small-RNA-mediated gene regulation.

## 5 Acknowledgement

Special thanks are due to Ms. Jia Fan of the Shenzhen X-institute for the introduction that enabled this academic collaboration. Code and data are available at https://github.com/cbaiming/DuplexFM.

## Footnotes

1 RNA-FM: https://huggingface.co/multimolecule/rnafm; mRNABERT: https://huggingface.co/YYLY66/mRNABERT.

## References

[1] David P Bartel. Micrornas: genomics, biogenesis, mechanism, and function. cell, 116(2):281–297, 2004.

[2] Benjamin P Lewis, Christopher B Burge, and David P Bartel. Conserved seed pairing, often flanked by adenosines, indicates that thousands of human genes are microrna targets. cell, 120(1):15–20, 2005.

[3] Sayda M Elbashir, Jens Harborth, Winfried Lendeckel, Abdullah Yalcin, Klaus Weber, and Thomas Tuschl. Duplexes of 21-nucleotide rnas mediate rna interference in cultured mammalian cells. nature, 411(6836):494–498, 2001.

[4] Victor Ambros. The functions of animal micrornas. Nature, 431(7006):350–355, 2004.

[5] Wigard P Kloosterman and Ronald HA Plasterk. The diverse functions of micrornas in animal development and disease. Developmental cell, 11(4):441–450, 2006.

[6] George A Calin and Carlo M Croce. Microrna signatures in human cancers. Nature reviews cancer, 6(11):857–866, 2006.

[7] David P Bartel. Micrornas: target recognition and regulatory functions. cell, 136(2):215–233, 2009.

[8] Hassan Dana, Ghanbar Mahmoodi Chalbatani, Habibollah Mahmoodzadeh, Rezvan Karimloo, Omid Rezaiean, Amirreza Moradzadeh, Narges Mehmandoost, Fateme Moazzen, Ali Mazraeh, Vahid Marmari, et al. Molecular mechanisms and biological functions of sirna. International journal of biomedical science: IJBS, 13(2):48, 2017.

[9] Lingxi Jiang, Yao Qi, Lei Yang, Yangbao Miao, Weiming Ren, Hongmei Liu, Yi Huang, Shan Huang, Shiyin Chen, Yi Shi, et al. Remodeling the tumor immune microenvironment via sirna therapy for precision cancer treatment. Asian Journal of Pharmaceutical Sciences, 18(5):100852, 2023.

[10] Richard W Carthew and Erik J Sontheimer. Origins and mechanisms of mirnas and sirnas. Cell, 136(4):642–655, 2009.

[11] Aimee L Jackson, Julja Burchard, Janell Schelter, B Nelson Chau, Michele Cleary, Lee Lim, and Peter S Linsley. Widespread sirna “off-target” transcript silencing mediated by seed region sequence complementarity. Rna, 12(7):1179–1187, 2006.

[12] Sergei A Manakov, Alexander A Shishkin, Brian A Yee, Kylie A Shen, Diana C Cox, Samuel S Park, Heather M Foster, Karen B Chapman, Gene W Yeo, and Eric L Van Nostrand. Scalable and deep profiling of mrna targets for individual micrornas with chimeric eclip. BioRxiv, pages 2022–02, 2022.

[13] Stephanie Sammut, Katarina Gresova, Dimosthenis Tzimotoudis, Eva Marsalkova, David Cechak, and Panagiotis Alexiou. mirbench: novel benchmark datasets for microrna binding site prediction that mitigate against prevalent microrna frequency class bias. Bioinformatics, 41(Supplement_1):i542–i551, 2025.

[14] Yilan Bai, Haochen Zhong, Taiwei Wang, and Zhi John Lu. Oligoformer: an accurate and robust prediction method for sirna design. Bioinformatics, 40(10):btae577, 2024.

[15] Anton Enright, Bino John, Ulrike Gaul, Thomas Tuschl, Chris Sander, and Debora Marks. Microrna targets in drosophila. Genome biology, 4(11):P8, 2003.

[16] Vikram Agarwal, George W Bell, Jin-Wu Nam, and David P Bartel. Predicting effective microrna target sites in mammalian mrnas. elife, 4:e05005, 2015.

[17] Giulia Riolo, Silvia Cantara, Carlotta Marzocchi, and Claudia Ricci. mirna targets: from prediction tools to experimental validation. Methods and protocols, 4(1):1, 2020.

[18] Caroline Diener, Andreas Keller, and Eckart Meese. The mirna–target interactions: an underestimated intricacy. Nucleic Acids Research, 52(4):1544–1557, 2024.

[19] Doron Betel, Anjali Koppal, Phaedra Agius, Chris Sander, and Christina Leslie. Comprehensive modeling of microrna targets predicts functional non-conserved and non-canonical sites. Genome biology, 11(8):R90, 2010.

[20] Ramkrishna Mitra and Sanghamitra Bandyopadhyay. Multimitar: a novel multi objective optimization based mirna-target prediction method. PloS one, 6(9):e24583, 2011.

[21] Jun Ding, Xiaoman Li, and Haiyan Hu. Tarpmir: a new approach for microrna target site prediction. Bioinformatics, 32(18):2768–2775, 2016.

[22] Albert Pla, Xiangfu Zhong, and Simon Rayner. miraw: A deep learning-based approach to predict microrna targets by analyzing whole microrna transcripts. PLoS computational biology, 14(7):e1006185, 2018.

[23] Tongjun Gu, Xiwu Zhao, William Bradley Barbazuk, and Ji-Hyun Lee. mitar: a hybrid deep learning-based approach for predicting mirna targets. BMC bioinformatics, 22(1):96, 2021.

[24] Seonwoo Min, Byunghan Lee, and Sungroh Yoon. Targetnet: functional microrna target prediction with deep neural networks. Bioinformatics, 38(3):671–677, 2022.

[25] Tingpeng Yang, Yu Wang, and Yonghong He. Tec-mitarget: enhancing microrna target prediction based on deep learning of ribonucleic acid sequences. BMC bioinformatics, 25(1):159, 2024.

[26] Baiming Chen. Leveraging disease association degree for high-accuracy microrna target prediction. eLife, 2026. Reviewed Preprint.

[27] Angela Reynolds, Devin Leake, Queta Boese, Stephen Scaringe, William S Marshall, and Anastasia Khvorova. Rational sirna design for rna interference. Nature biotechnology, 22(3):326–330, 2004.

[28] Kumiko Ui-Tei, Yuki Naito, Fumitaka Takahashi, Takeshi Haraguchi, Hiroko Ohki-Hamazaki, Aya Juni, Ryu Ueda, and Kaoru Saigo. Guidelines for the selection of highly effective sirna sequences for mammalian and chick rna interference. Nucleic acids research, 32(3):936–948, 2004.

[29] Yuki Naito, Tomoyuki Yamada, Kumiko Ui-Tei, Shinichi Morishita, and Kaoru Saigo. sidirect: highly effective, target-specific sirna design software for mammalian rna interference. Nucleic acids research, 32(suppl_2):W124–W129, 2004.

[30] Zhi John Lu and David H Mathews. Oligowalk: an online sirna design tool utilizing hybridization thermodynamics. Nucleic acids research, 36(suppl_2):W104–W108, 2008.

[31] Dieter Huesken, Joerg Lange, Craig Mickanin, Jan Weiler, Fred Asselbergs, Justin Warner, Brian Meloon, Sharon Engel, Avi Rosenberg, Dalia Cohen, et al. Design of a genome-wide sirna library using an artificial neural network. Nature biotechnology, 23(8):995–1001, 2005.

[32] Jean-Philippe Vert, Nicolas Foveau, Christian Lajaunie, and Yves Vandenbrouck. An accurate and interpretable model for sirna efficacy prediction. BMC bioinformatics, 7(1):520, 2006.

[33] Ryan L Boudreau, Ryan M Spengler, Ray H Hylock, Brandyn J Kusenda, Heather A Davis, David A Eichmann, and Beverly L Davidson. sispotr: a tool for designing highly specific and potent sirnas for human and mouse. Nucleic acids research, 41(1):e9–e9, 2013.

[34] Hugo Dalla-Torre, Liam Gonzalez, Javier Mendoza-Revilla, Nicolas Lopez Carranza, Adam Henryk Grzywaczewski, Francesco Oteri, Christian Dallago, Evan Trop, Bernardo P De Almeida, Hassan Sirelkhatim, et al. Nucleotide transformer: building and evaluating robust foundation models for human genomics. Nature Methods, 22(2):287–297, 2025.

[35] Jiayang Chen, Zhihang Hu, Siqi Sun, Qingxiong Tan, Yixuan Wang, Qinze Yu, Licheng Zong, Liang Hong, Jin Xiao, Tao Shen, et al. Interpretable rna foundation model from unannotated data for highly accurate rna structure and function predictions. arXiv preprint arXiv:2204.00300, 2022.

[36] Ning Wang, Jiang Bian, Yuchen Li, Xuhong Li, Shahid Mumtaz, Linghe Kong, and Haoyi Xiong. Multi-purpose rna language modelling with motif-aware pretraining and type-guided fine-tuning. Nature Machine Intelligence, 6(5):548–557, 2024.

[37] Rafael Josip Penić, Tin Vlašić, Roland G Huber, Yue Wan, and Mile Šikić. Rinalmo: General-purpose rna language models can generalize well on structure prediction tasks. Nature Communications, 16(1):5671, 2025.

[38] Alexander Rives, Joshua Meier, Tom Sercu, Siddharth Goyal, Zeming Lin, Jason Liu, Demi Guo, Myle Ott, C Lawrence Zitnick, Jerry Ma, et al. Biological structure and function emerge from scaling unsupervised learning to 250 million protein sequences. Proceedings of the national academy of sciences, 118(15):e2016239118, 2021.

[39] Ying Xiong, Aowen Wang, Yu Kang, Chao Shen, Chang-Yu Hsieh, and Tingjun Hou. mrnabert: advancing mrna sequence design with a universal language model and comprehensive dataset. Nature Communications, 16(1):10371, 2025.

[40] Eva Klimentová, Václav Hejret, Ján Krčmář, Katarína Grešová, Ilektra-Chara Giassa, and Panagiotis Alexiou. mirbind: A deep learning method for mirna binding classification. Genes, 13(12):2323, 2022.

[41] Vaclav Hejret, Nandan Mysore Varadarajan, Eva Klimentova, Katarina Gresova, Ilektra-Chara Giassa, Stepanka Vanacova, and Panagiotis Alexiou. Analysis of chimeric reads characterises the diverse targetome of ago2-mediated regulation. Scientific Reports, 13(1):22895, 2023.

[42] Eric L Van Nostrand, Gabriel A Pratt, Alexander A Shishkin, Chelsea Gelboin-Burkhart, Mark Y Fang, Balaji Sundararaman, Steven M Blue, Thai B Nguyen, Christine Surka, Keri Elkins, et al. Robust transcriptome-wide discovery of rna-binding protein binding sites with enhanced clip (eclip). Nature methods, 13(6):508–514, 2016.

[43] Aleksandra Helwak, Grzegorz Kudla, Tatiana Dudnakova, and David Tollervey. Mapping the human mirna interactome by clash reveals frequent noncanonical binding. Cell, 153(3):654–665, 2013.

[44] Takayuki Katoh and Tsutomu Suzuki. Specific residues at every third position of sirna shape its efficient rnai activity. Nucleic acids research, 35(4):e27, 2007.

[45] Mohammed Amarzguioui, Torgeir Holen, Eshrat Babaie, and Hans Prydz. Tolerance for mutations and chemical modifications in a sirna. Nucleic acids research, 31(2):589–595, 2003.

[46] Jens Harborth, Sayda M Elbashir, Kim Vandenburgh, Heiko Manninga, Stephen A Scaringe, Klaus Weber, and Thomas Tuschl. Sequence, chemical, and structural variation of small interfering rnas and short hairpin rnas and the effect on mammalian gene silencing. Antisense and Nucleic Acid Drug Development, 13(2):83–105, 2003.

[47] Andrew C Hsieh, Ronghai Bo, Judith Manola, Francisca Vazquez, Olivia Bare, Anastasia Khvorova, Stephen Scaringe, and William R Sellers. A library of sirna duplexes targeting the phosphoinositide 3-kinase pathway: determinants of gene silencing for use in cell-based screens. Nucleic acids research, 32(3):893–901, 2004.

[48] Timothy A Vickers, Seongjoon Koo, C Frank Bennett, Stanley T Crooke, Nicholas M Dean, and Brenda F Baker. Efficient reduction of target rnas by small interfering rna and rnase h-dependent antisense agents: a comparative analysis. Journal of Biological Chemistry, 278(9):7108–7118, 2003.

[49] Xiaowei Wang, Xiaohui Wang, Rajeev K Varma, Lesslie Beauchamp, Susan Magdaleno, and Timothy J Sendera. Selection of hyperfunctional sirnas with improved potency and specificity. Nucleic acids research, 37(22):e152–e152, 2009.

[50] Wenchong Tan, Mingshu Dai, Shimin Ye, Xin Tang, Dawei Jiang, Dong Chen, and Hongli Du. Ensirna: a multimodality method for sirna-mrna and modified sirna efficacy prediction based on geometric graph neural network. Journal of Molecular Biology, 437(12):169131, 2025.

[51] Lei Sun, Kui Xu, Wenze Huang, Yucheng T Yang, Pan Li, Lei Tang, Tuanlin Xiong, and Qiangfeng Cliff Zhang. Predicting dynamic cellular protein–rna interactions by deep learning using in vivo rna structures. Cell research, 31(5):495–516, 2021.

[52] Robert C Spitale, Ryan A Flynn, Qiangfeng Cliff Zhang, Pete Crisalli, Byron Lee, Jong-Wha Jung, Hannes Y Kuchelmeister, Pedro J Batista, Eduardo A Torre, Eric T Kool, et al. Structural imprints in vivo decode rna regulatory mechanisms. Nature, 519(7544):486–490, 2015.

[53] Martin Mann, Patrick R Wright, and Rolf Backofen. Intarna 2.0: enhanced and customizable prediction of rna–rna interactions. Nucleic acids research, 45(W1):W435–W439, 2017.

[54] Stefan Ludwig Ameres, Javier Martinez, and Renée Schroeder. Molecular basis for target rna recognition and cleavage by human risc. Cell, 130(1):101–112, 2007.

[55] Michael Kertesz, Nicola Iovino, Ulrich Unnerstall, Ulrike Gaul, and Eran Segal. The role of site accessibility in microrna target recognition. Nature genetics, 39(10):1278–1284, 2007.

[56] Edward J Hu, Yelong Shen, Phillip Wallis, Zeyuan Allen-Zhu, Yuanzhi Li, Shean Wang, Liang Wang, Weizhu Chen, et al. Lora: Low-rank adaptation of large language models. Iclr, 1(2):3, 2022.

